# Optimization of an anti *Staphylococcus* antibiotic produced by tropical soil dwelling *Streptomyces parvulus*

**DOI:** 10.1101/060392

**Authors:** Sonashia Velho-Pereira, Nandkumar M Kamat

## Abstract

An antibiotic produced by strain *Streptomyces parvulus* showing activity against *Staphylococcus citreus* was subjected to various optimization parameters for enhancing its production. Nutritional and physiological parameters produced by *S. parvulus* under shaken flask conditions were determined. Optimization of these parameters led to 11% increase in antibiotic activity with a mean zone of inhibition of 42 mm.

Highest antibiotic production was obtained at 250 rpm for 14 days with optimum temperature of 28°C and pH 7. Kuster#x2019;s modified medium containing glycerol 0.7% (v/v), casein 0.03% (w/v), NaCl 0% (w/v), phosphate 0.25% (w/v), KNO3 0.1% (w/v) and CaCO3 0.0015% (w/v) concentration was found ideal.

## 1. Introduction

Novel antibiotics are continuously in demand due to the inevitable rise of antibiotic-resistant strains of pathogenic bacteria, reducing morbidity and mortality of life’ s expectancy (Fischbach and Walsh 2009). Among the pathogenic bacteria *Staphylococcus citreus* needs a special mention on account of its pathogenicity.

*S. citreus*, a virulent strain has risen due to spontaneous split off of pure line strains of pathogenic *S. aureus* (Pinner and Voldrich 1932) which causes skin infections, pneumonia, meningitis, endocarditis, toxic shock syndrome, septicemia and arthritis (Dilsen et al. 1961; Chambers 2001).

Research done to harness potential drug candidates against this pathogen has not been reported, although concerted efforts to harness potential drugs against multi drug resistant strains of *Staphylococcus aureus* such as MRSA is underway (Demain and Sanchez 2009).

Actinobacteria have proven to be prolific producers of secondary metabolites among all microbial organisms, accounting for 45% of all microbial metabolites of which 80% (7,600 compounds) are produced by genus *Streptomyces* (Berdy 2005).

According to Watve et al. (2001) predictive modeling of genus *Streptomyces* suggests that over 150,000 bioactive metabolites from this genus still needs to be discovered.

Antibiotic biosynthesis in *Streptomycetes* has been reported to be highly dependent on the nutritional and physiological factors prevailing during its growth as it helps in cell proliferation which expresses genetic information favoring secondary metabolism (Abbanat et al. 1999). These metabolic processes are species specific and can be enhanced or minimized under different physiological conditions (Yarbrough et al. 1992; Abbanat et al. 1999; Selvin et al. 2009; Visalakchi and Muthumary 2009; Rezuanul et al. 2009; Elleuch et al. 2010; Nanjwade et al. 2010; Panda et al. 2011; Chhabra and Keasling 2011; Darabpour et al. 2012; Mangamuri et al. 2012; Singh and Rai 2012; Luthra and Dubey 2012; Gunda and Charya 2013). Thus, it is essential to standardize growth conditions of the producer strain, for maximum synthesis of its bioactive molecules (Sujatha et al. 2005; Olmos et al. 2013).

Classical strain improvement despite being laborious and time consuming is still widely used due to its high success rate behind improved production titers of antibiotics such as penicillin, cephalosporin C, tylosin, salinomycin, chlortetracycline and tetracycline (Chhabra and Keasling 2011).

In the course of screening actinobacteria for antibiotic compounds, a strain identified as *Streptomyces parvulus* showed broad spectrum activity as revealed by perpendicular streak (Badji et al. 2007) and agar well diffusion method (Devillers et al. 1989). The present communication deals with the optimal parameters required for maximizing antibiotic production by *S. parvulus* against pathogenic bacteria *Staphylococcus citreus*: We claim this to be the first such report.

## 2. Methods

### 2.1. Location and sampling

The actinobacterial strain CFA-9, deposited at Goa University Fungal Culture Collection (GUFCC 20101) was isolated from forest soil, Canacona, Goa, India (latitude 14° 59’45.76”N and longitude 74° 03’02.17”E). Isolation of this strain was carried out using a novel baiting technique, which employs a microcosm with Arginine Vitamin Agar (AVA) medium coated slides to specifically capture *ex situ* actinobacterial diversity (Velho-Pereira and Kamat 2011; 2012).

### 2.2. Taxonomic and molecular identification of the producer organism

The morphological and cultural characteristics of the strain CFA-9 were studied by using traditional criteria of classification (Locci 1989; Cross and Goodfellow 1973). The micromorphological studies were done using light and scanning electron microscopy (SEM) (Williams and Davies 1967).

For molecular identification, genomic DNA of the strain was extracted and its quality was evaluated as a single distinct band on 1.2% agarose gel. The fragment 16S rRNA gene was amplified by PCR and the amplicon was purified to remove contaminants. Forward and reverse DNA sequencing reaction of PCR amplicon was carried out with 8F and 1 492R primers using BDT v3.1 cycle sequencing kit on ABI 3730×l Genetic analyser. Consensus sequence of 1347bp 16S rRNA gene was generated from forward and reverse sequence data using aligner software. This work was done at Xcelris Labs Ltd. (www.xcelerislabs.com). The sequence has been deposited in Genebank database (http://www.ncbi.nlm.nih.gov/genbank/submit).

Phylogenetic analyses were conducted using MEGA v5 (Tamura et al. 2011). The 16S rRNA gene sequence of strain CFA-9 was aligned using the Clustal W program against corresponding nucleotide sequences of representatives of *Streptomyces* genus retrieved from Genbank (http://www.ncbi.nlm.nih.gov/genbank). Phylogenetic tree was inferred by the maximum-likelihood method (Felsenstein 1985) based on the Hasegawa-Kishino-Yano (HKY) (Hasegawa et al. 1985) model. Tree topologies were evaluated by bootstrap analysis (Felsenstein 1985) based on 1000 resamplings.

### 2.3. Antibiotic production medium

Kuster’s broth (Kuster and William 1964) composed of glycerol (0.8% v/v), Casein (0.03% w/v), NaCl (2% w/v), KNO_3_ (0.2 w/v), K_2_HPO_4_ (0.2% w/v), MgSO4.7H2O (0.005% w/v), CaCO_3_ (0.001% w/v), pH 7 was used as the basic fermentation medium. It was inoculated with a seven day old culture (5% inoculum) under sterile conditions. The culture flasks were fixed on a rotary shaker (Orbitek^R^, Scigenics Biotech, Pvt. Ltd., India) 250 rpm; rotation diameter: 2.0 cm, placed in a thermostated cabinet at 28°C for fourteen days fermentation process. All these parameters formed the positive control of the experiment. The chemicals were procured from HiMedia, Mumbai, India.

### 2.4. Antibiotic bioassay and test organism

Agar well diffusion method (Devillers et al. 1989) was used for detection of antimicrobial activity. Antibiotic bioassay was carried out using the cell free centrifugate (CFC) to detect extracellular production of bioactive metabolites. Uninoculated Kuster’s broth was used as a negative control.

The culture broth was subjected to centrifugation at 4000 g for 20 min to obtain a CFC. Medium for bioassay was Muller Hinton agar (Himedia veg, Mumbai, India). Three cores of 6 mm diameter were excised from the Mueller Hinton agar plates, pre-seeded with the test organism using sterile swabs. The wells were filled with the supernatant (50-70 μl) using Accupipet model T1000 (Tarsons Products Pvt. Ltd., Kolkata, India).The plates were incubated at 28°C for 48h and inhibition zones (ZOI) were visualized and measured in millimeters.

Since preliminary screening showed highest activity against *Staphylococcus citreus*, the same was chosen as a test pathogenic strain for optimizing all parameters. The test organism *S. citreus* was procured from Department of Microbiology, Goa Medical College, Goa and maintained on nutrient agar media (Himedia, Mumbai, India).

### 2.5. Optimization parameters

Antibiotic production was optimized by using different physiological and nutritional parameters *viz*., days of incubation, temperature, pH, sodium chloride, organic and inorganic carbon and nitrogen sources and phosphate. The optimum conditions identified for one parameter were used for optimizing the other parameters sequentially. The efficiency of the optimized parameters was established on the basis of zone of inhibition (ZOI) in mm. All experiments were performed in triplicates (n = 3).

#### 2.5.1. Days of incubation

To study the effect of incubation period on antibiotic production,1 0 ml aliquot of the culture broth was collected aseptically at regular intervals of 4, 7, 1 2, 1 4, 1 6, 1 8 and 20^th^ days and the CFC was subjected to bioassay.

#### 2.5.2. Temperature

The optimum temperature for antibiotic production was assayed by incubating the production medium at 25, 28, 32 and 37°C. The control was maintained at 28°C.

#### 2.5.3. pH

Influence of pH on antibiotic production of the strain was determined by adjusting the pH of production medium ranging from 3 - 11 with 0.1 N NaOH/0.1 N HCl. pH 7 was used as the control.

#### 2.5.4. NaCl concentration (ppm)

The effect of sodium chloride on antibiotic production was studied using different salinity concentrations of 0; 1 0,000; 1 5,000; 20,000; 25,000; 30,000 and 35,000 ppm. NaCl concentration of 20,000 ppm was kept as control.

#### 2.5.5 Organic Carbon and Nitrogen source(%)

To study the influence of carbon and nitrogen source on antibiotic production, varied concentrations of the respective sources were tested. Glycerol being the sole organic carbon source of the Kuster’s broth medium was studied using its varied concentration of 0.5, 0.6, 0.7, 0.8, 0.9, 1, 1.2 and 1.3% (v/v). Glycerol concentration of 0.8% (v/v) was kept as control. Casein being the sole organic nitrogen source of the medium was studied using its varied concentration of 0.01-0.09% (w/v). Casein concentration of 0.03% (w/v) was kept as control.

#### 2.5.6. Phosphate concentration (%)

To study the effect of phosphate mineral (K2HPO4) on antibiotic production different concentrations, 0.00029; 0.00057; 0.00086; 0.0011; 0.0014; 0.0017; 0.002; 0.0023; 0.0026 and 0.0029% (w/v) were tested. Phosphate concentration of 0.0011% (w/v) was kept as control.

#### 2.5.7. Inorganic nitrogen and carbon source (%)

The influence of inorganic nitrogen (KNO3) and carbon source (CaCO3) present in the antibiotic production medium was studied using varied KNO_3_ concentration of 0.1, 0.15, 0.2, 0.25, 0.3, 0.35, 0.4, 0.45, 0.5 and 0.55% (w/v). KNO_3_ concentration of 0.1% (w/v) was kept as control and eight varied concentration of CaCO_3_ i.e. 0, 0.0005, 0.001, 0.0015, 0.002, 0.0025, 0.003, 0.0035% (w/v) was studied. CaCO_3_ concentration of 0.001% (w/v) was kept as control.

### 2.6. Statistical Analysis

Results are expressed as a mean of three experiments ± standard deviations (SDs). Statistical analyses were performed using WASP (Web Based Agricultural Statistics Software Package 1.0, (http://www.icargoa.res.in/wasp/index.php) and differences were considered significant if p ≤ 0.05.

## 3. Results

### 3.1. Selection and molecular identification of the strain

Among the five strains isolated from the forest soil, CFA-9 with broad spectrum antimicrobial activity of 66.7% against human pathogens namely Gram negative bacteria *Shigella flexneri*, *Enterobacter aerogens* and Gram positive bacteria *Bacillus subtilis*, *Staphylococcus typhi* and *S. citreus* was selected. Highest zone of inhibition (31 mm) was observed against *S. citreus*. The strain exhibited grey aerial mycelium, with spiral spore chains and produced a bright yellow pigment and was identified as *Streptomyces parvulus*, with gene bank accession number of KC904376 (Fig. 1).

**Fig 1.**
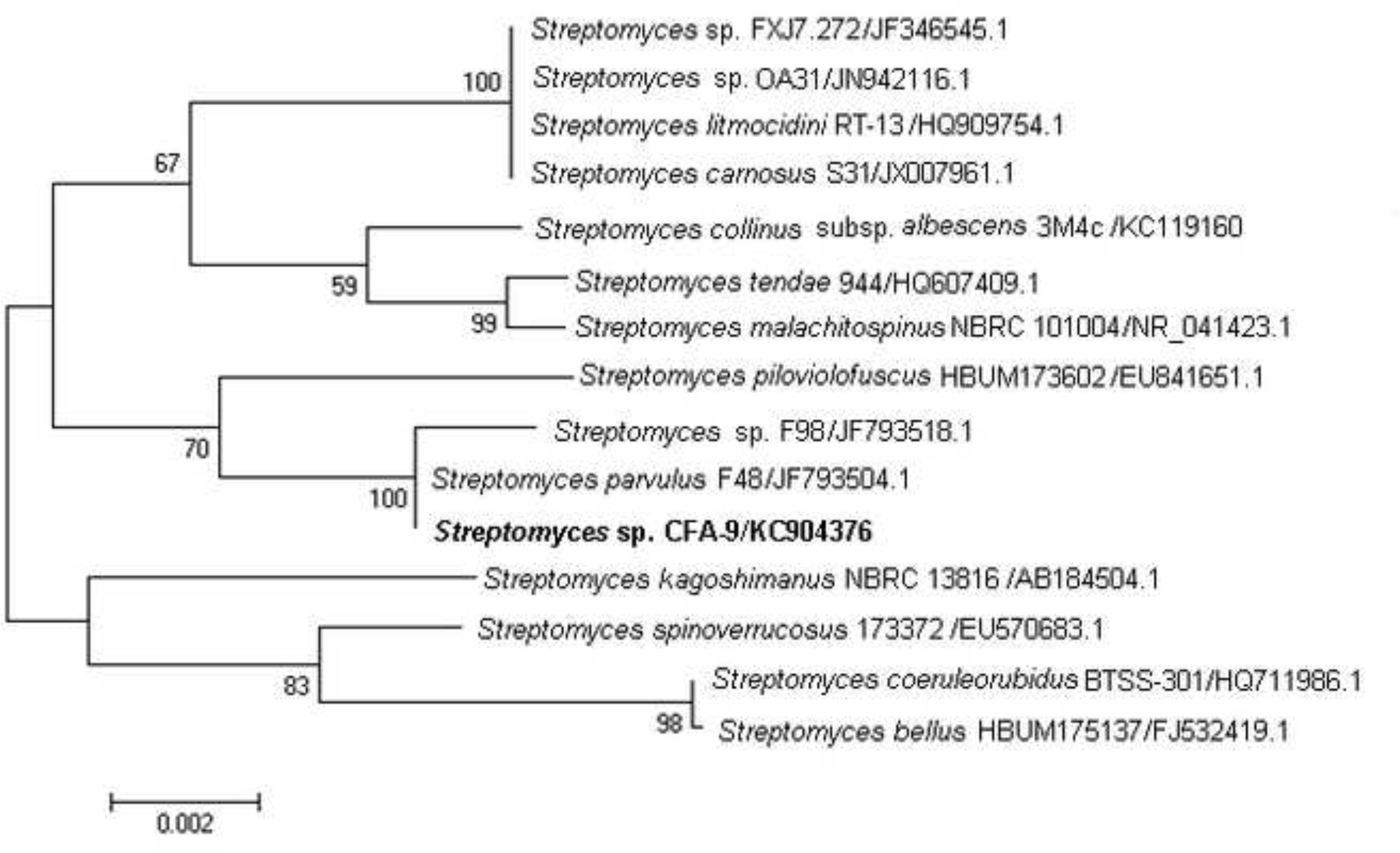
Maximum-likelihood tree based on 16S rRNA gene sequence showing the relations between antibiotic producer strain *Streptomyces* strain CFA-9 and type species of the genus *Streptomyces* within the order actinomycetales under actinobacteria. The numbers at the nodes indicate the levels of bootstrap support based on maximum-likelihood analyses of 1000 resampled data sets (only values >50% are shown). The scale bar indicates 0.002 substitutions per nucleotide positions.

### 3.2. Optimization parameters

#### 3.2.1. Days of incubation

The maximum antibiotic production resulting in a mean ZOI of 35 mm was recorded over a period of 14 days. The activity was observed from the 4^th^ day of incubation and reached a maximum on the 14^th^ day showing significant variation (p ≤ 0.05). Thereafter, with the increase in incubation period, the antibiotic production decreased (Fig. 2).

**Fig 2.**
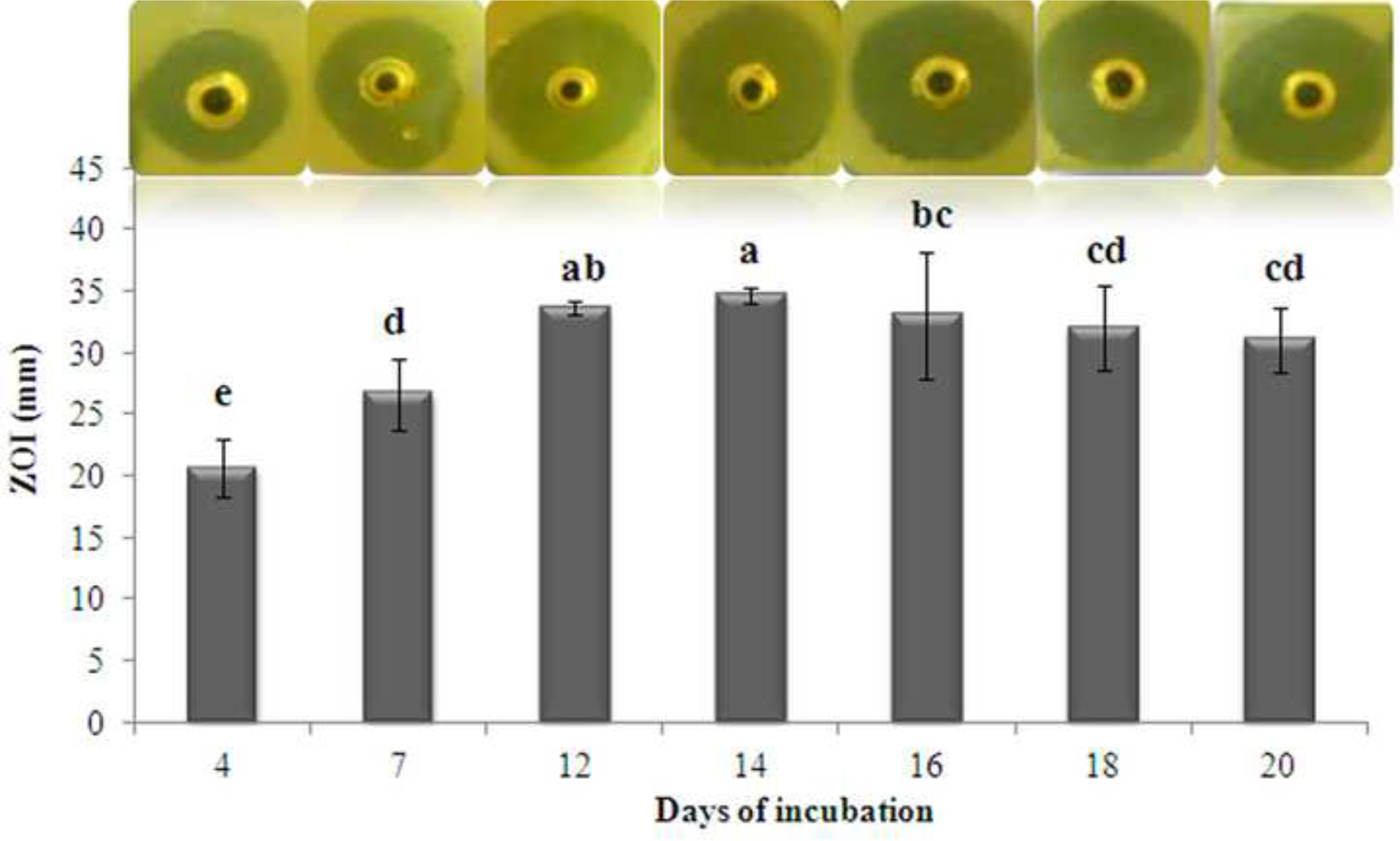
Effect of incubation periods on antibiotic production. Bars indicate standard devi ati on of the mean and the superscripts indicate significant differences (p ≤ 0.05).

#### 3.2.2. Temperature

As seen in the Fig. 3, incremental temperature rise led to an increase in the antibiotic production till it reached the optimum, further increase in temperature was accompanied by a decrease in the antibiotic production. Maximum yield of bioactive metabolites was observed when the strain was cultured at optimum temperature of 28°C with a mean ZOI of 37 mm which was significant (p ≤ 0.05). 25°C and 32°C also showed an appreciable mean ZOI of 29 and 32 mm respectively. The lowest mean ZOI of 24 mm was observed at 37^o^C (Fig. 3).

**Fig 3.**
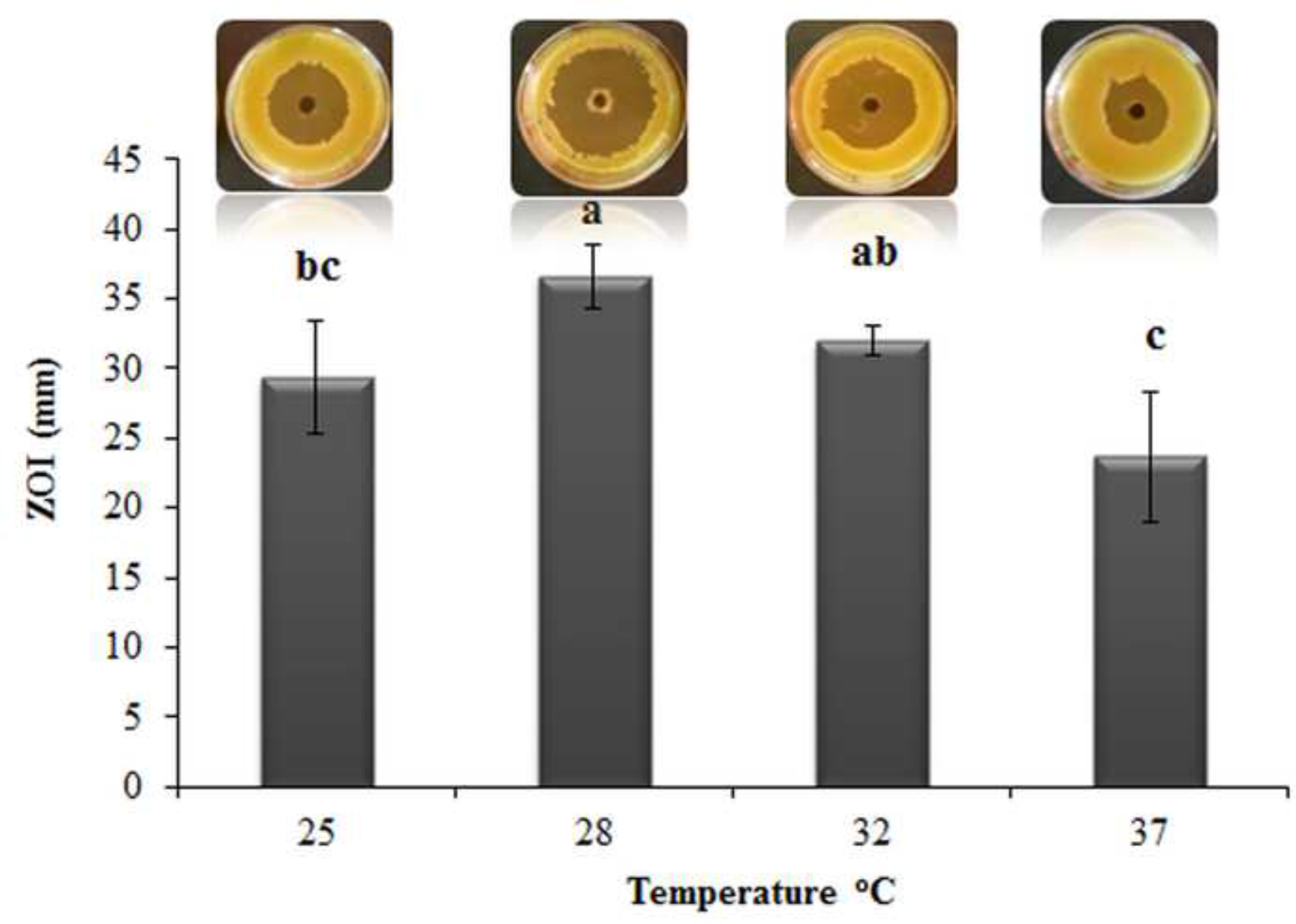
Effect of temperature on antibiotic production. Bars indicate standard devi ati on of the mean and the superscripts indicate significant differences (p ≤ 0.05).

#### 3.2.3. pH

Maximum antibiotic activity occurred at pH 7.0, exhibiting mean ZOI of 37 mm which was significant (p ≤ 0.05). No antibiotic production was observed at pH 3, 4 and 11. Increasing the pH value led to an increase in the antibiotic production up to certain threshold limit and further increase in values resulted in decrease in the antibiotic production (Fig. 4).

**Fig 4.**
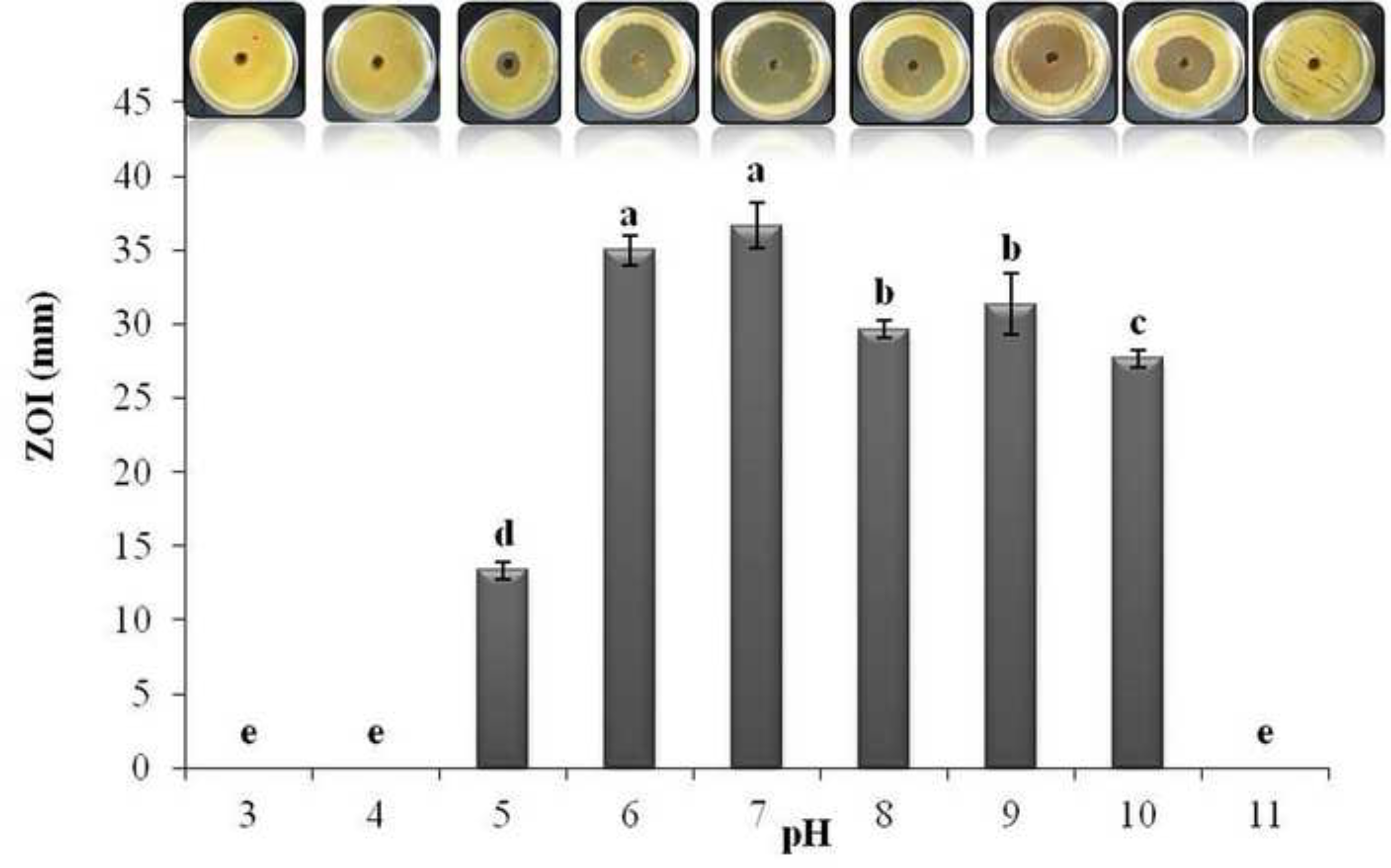
Effect of pH on antibiotic production. Bars indicate standard deviation of the mean and the superscripts indicate significant differences (p ≤ 0.05).

#### 3.2.4. NaCl concentration

Maximum antibiotic production in terms of mean ZOI of 37 mm, was obtained without NaCl and was significant (p ≤ 0.05). A significant difference was found over control (20,000 ppm concentration) exhibiting a mean ZOI of 33 mm. However activity was observed at all concentration of NaCl (Fig. 5).

**Fig 5.**
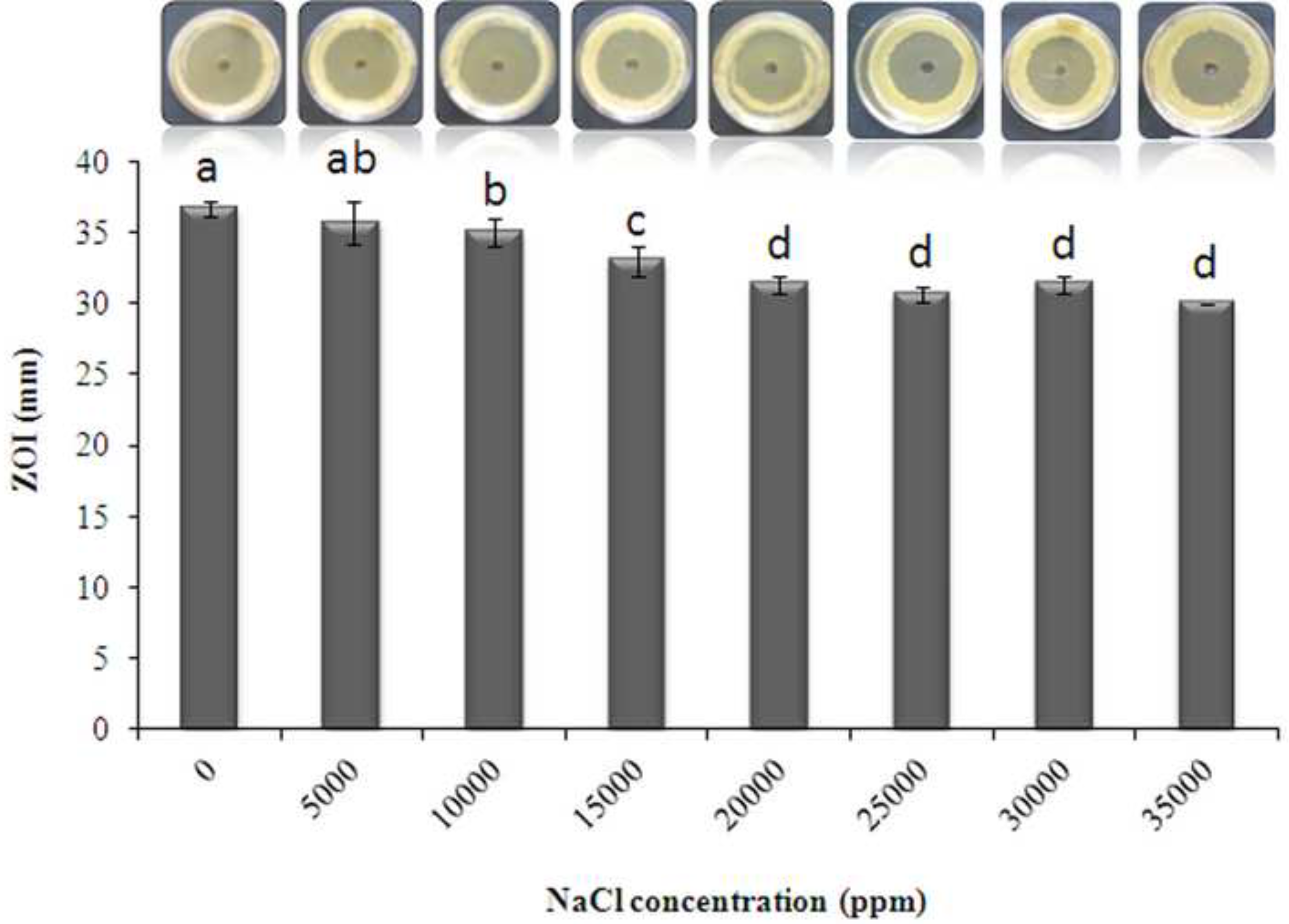
Effect of NaCl (ppm) on antibiotic production. Bars indicate standard devi ati on of the mean and the superscripts indicate significant differences (p ≤ 0.05).

#### 3.2.4. Organic Carbon and Nitrogen concentration (%)

The onset and intensity of secondary metabolism is dependent on various nutritional factors like carbon and nitrogen sources. Maximum antibiotic activity was observed with 0.7% glycerol concentration showing a mean ZOI of 35 mm and was significant (p ≤ 0.05). Data indicated that increasing concentration of glycerol from 0.5-0.7% led to an increase in the antibiotic production, thereafter further increase in values from 0.8-1.2% resulted in its decrease (Fig. 6).

**Fig 6.**
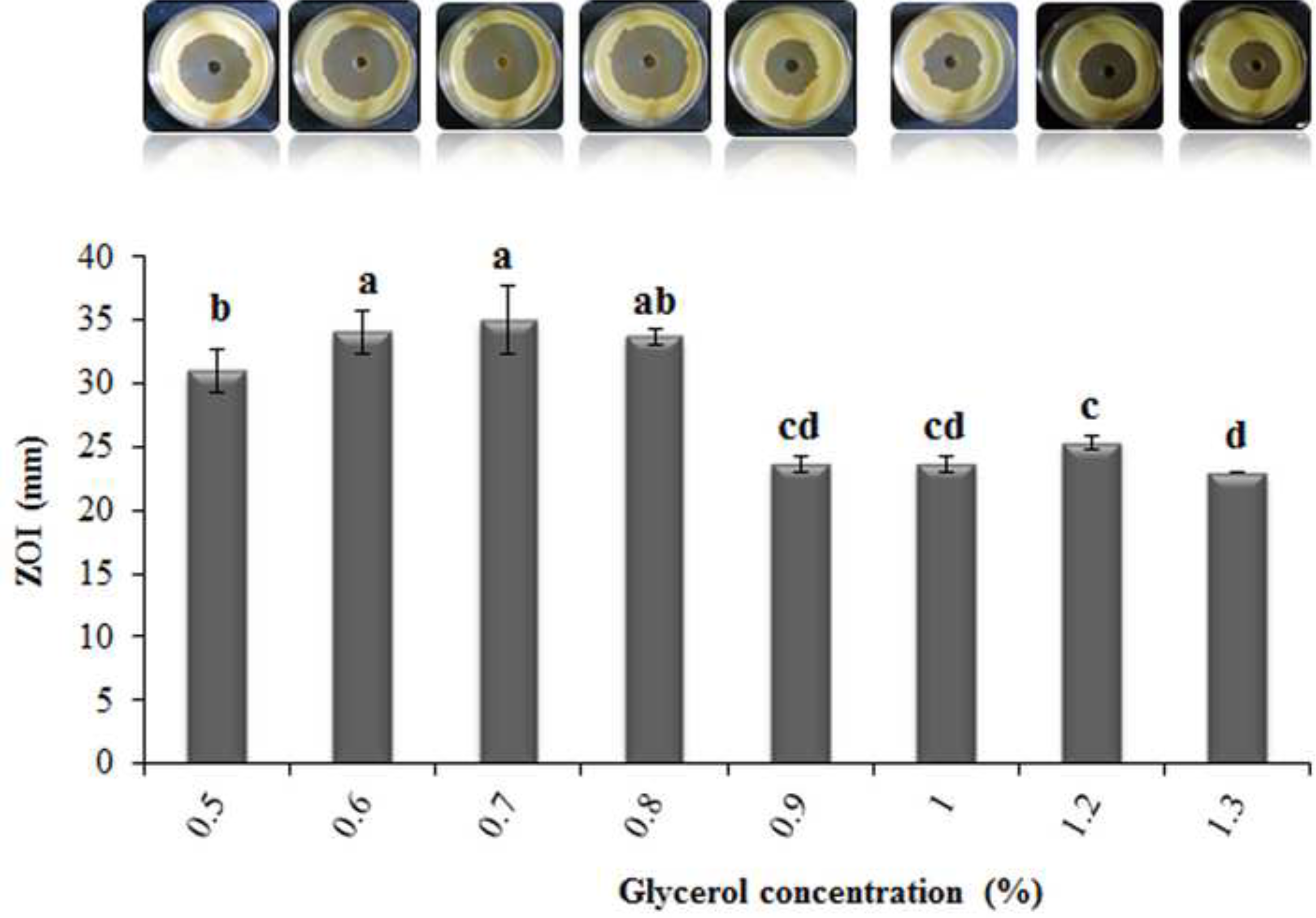
Effect of different concentrations of glycerol (%) as organic carbon source on antibiotic production. Bars indicate standard deviation of the mean and the superscripts indicate significant differences (p ≤ 0.05).

Maximum antibiotic activity was observed with 0.03% casein concentration showing a mean ZOI of 39 mm and was significant (p ≤ 0.05). However, other concentration of casein also favoured the production of antibiotic compounds (Fig. 7).

**Fig 7.**
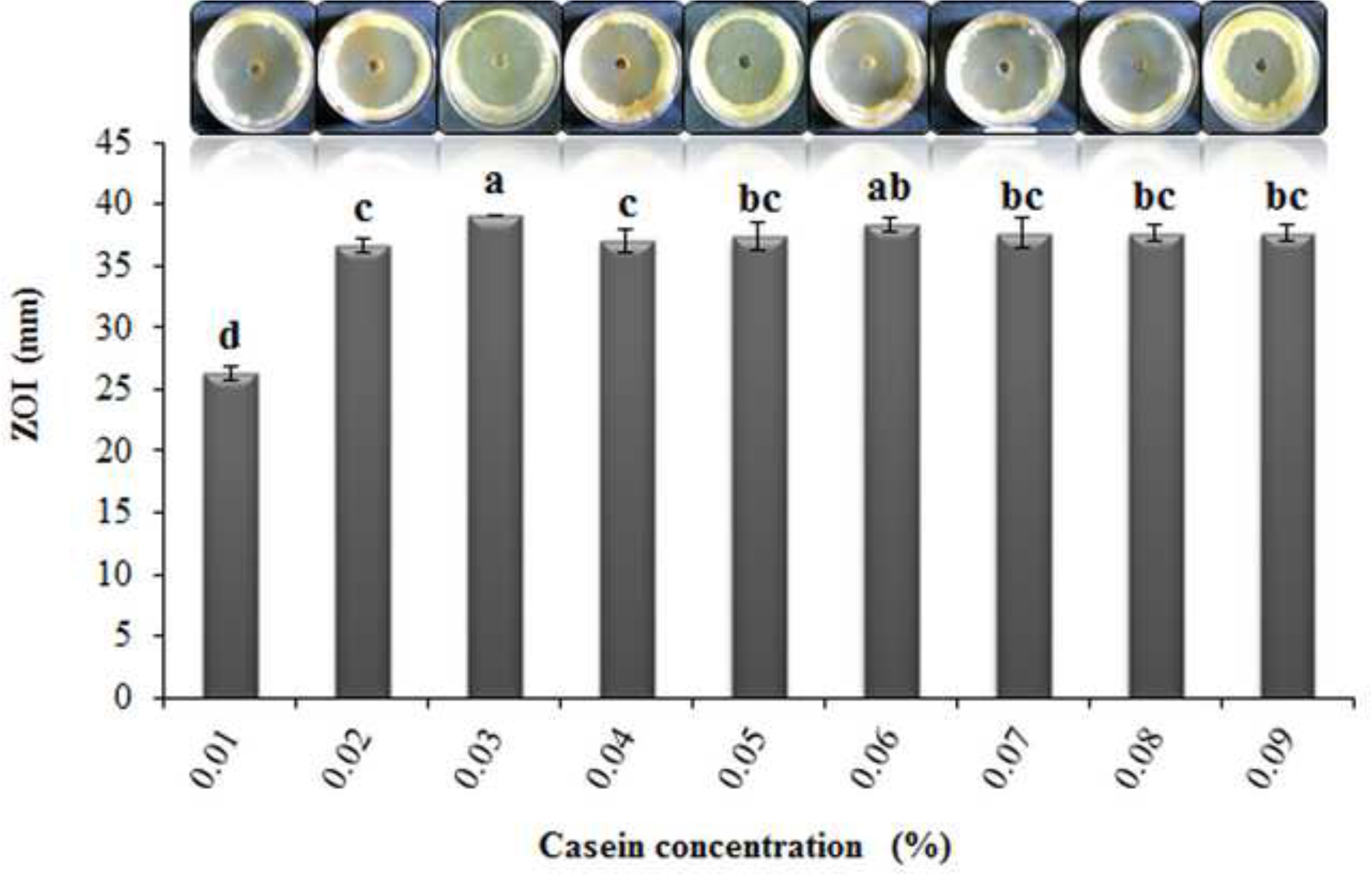
Effect of different concentrations of casein (%) as organic nitrogen source on antibiotic production. Bars indicate standard deviation of the mean and the superscripts indicate significant differences (p ≤ 0.05).

#### 3.2.5. Phosphate concentration (%)

As seen in Fig. 9, 0.25% (w/v) concentration of phosphate resulted in maximum yield of antibiotic exhibiting a mean ZOI of 35 mm and was significant (p ≤ 0.05). The results also indicated other concentrations of phosphate exhibiting an appreciable ZOI (Fig. 8).

**Fig 8.**
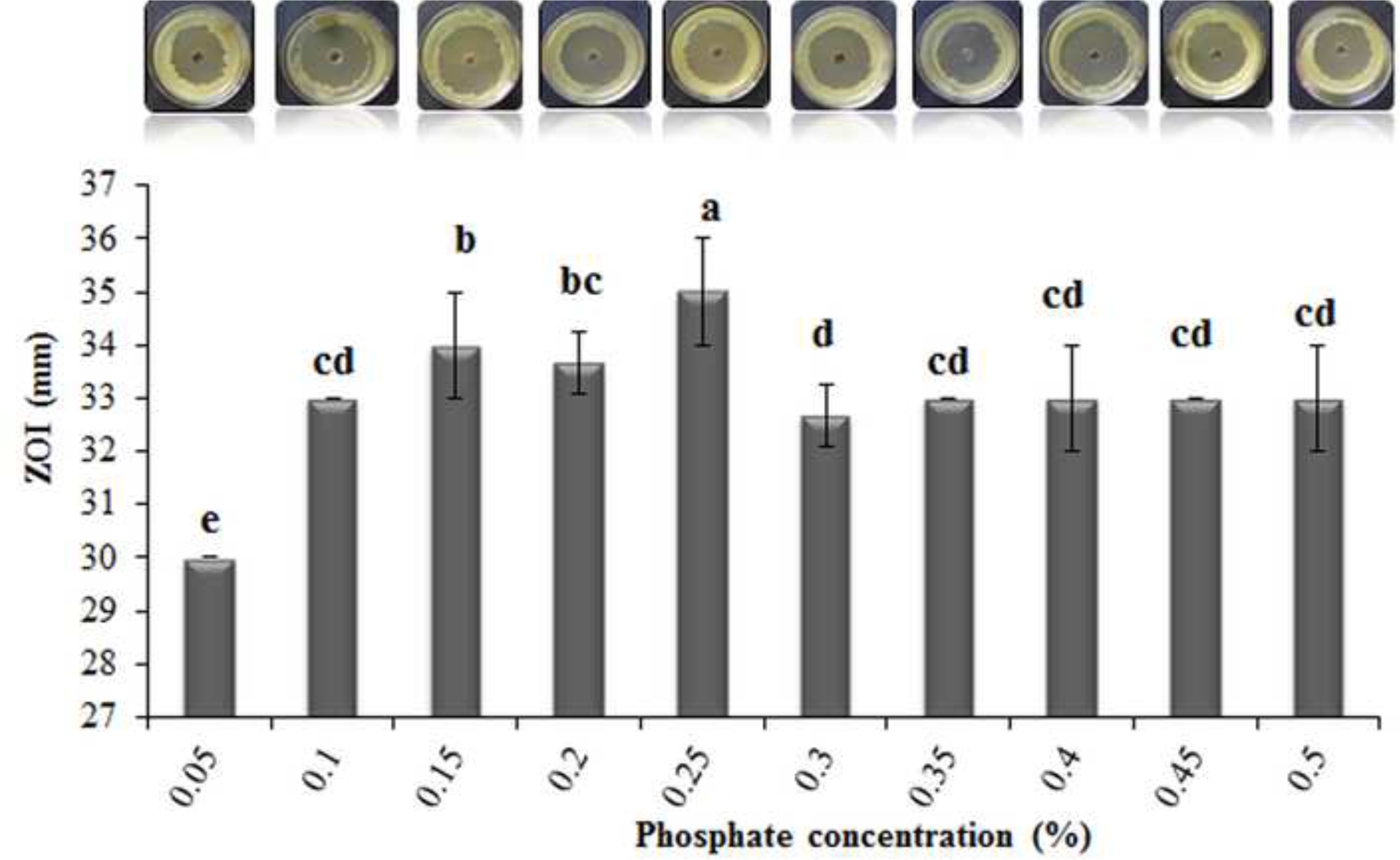
Effect of phosphate (%) on antibiotic production. Bars indicate standard devi ati on of the mean and the superscripts indicate significant differences (p ≤ 0.05).

**Fig 9.**
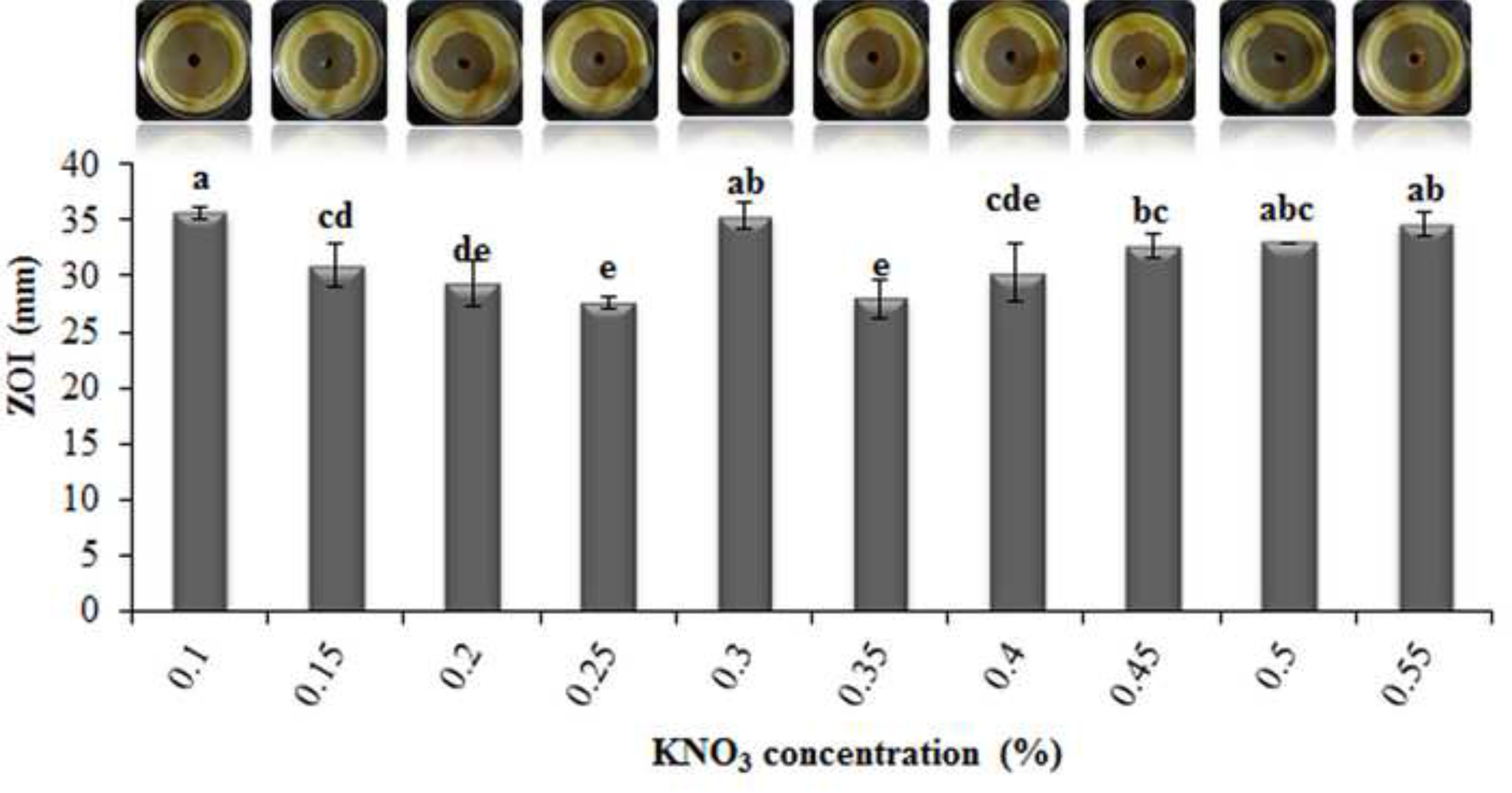
Effect of different concentrations of KNO_3_ (%) as inorganic nitrogen source on antibiotic production. Bars indicate standard deviation of the mean and the superscripts indicate significant differences (p ɤ 0.05).

#### 3.2.6. Inorganic Nitrogen and Carbon concentration (%)

Maximum antibiotic activity was observed with its 0.1% KNO_3_ concentration exhibiting a mean ZOI of 36 mm which was much higher to that of control (0.2%) with a mean ZOI of 29 mm and was significant (p ≤ 0.05) (Fig. 9). Maximum antibiotic activity was observed with 0.0015% and 0.002% CaCO_3_ concentration with a mean ZOI of 37 mm in both cases and was significant (p ≤ 0.05) (Fig. 10).

**Fig 10.**
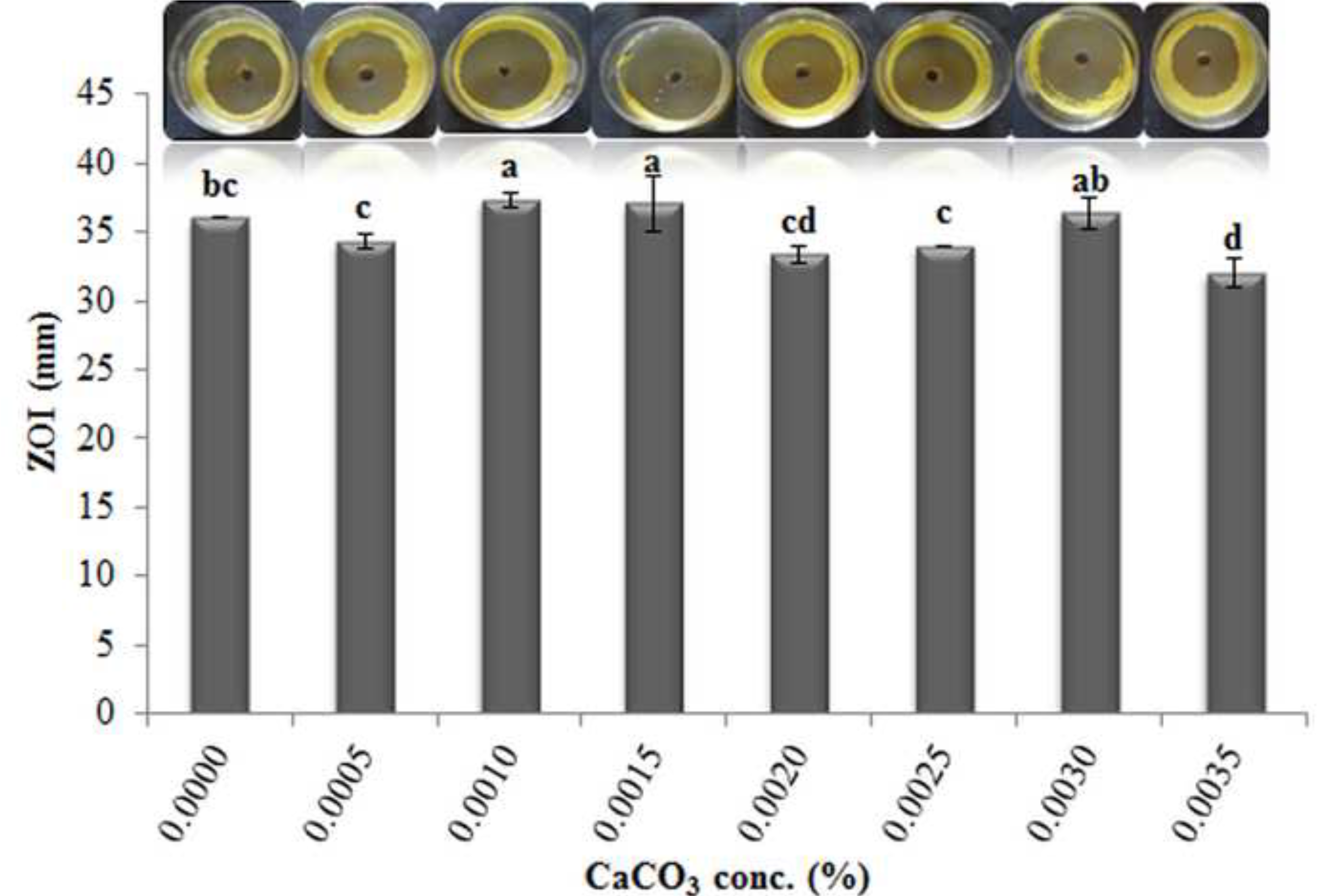
Effect of different concentrations of CaCO_3_ (%) as inorganic carbon source on antibiotic production. Bars indicate standard deviation of the mean and the superscripts indicate signific ant differences (p ≤ 0.05).

## 4. Discussion

A broad-spectrum antibiotic producing strain *Streptomyces parvulus* exhibiting maximum activity against human pathogenic bacteria *Staphylococcus citreus* was subjected to various optimization parameters.

The ability of actinobacterial cultures to produce antibiotics is not fixed and arises from intracellular intermediates through defined biochemical pathways. It can either be greatly increased or completely lost depending on the conditions in which they are grown (Kavitha and Vijayalakshmi 2009). The production of most antibiotics is regulated by complex biosynthetic pathways encoded by physically clustered genes (Sevcikova and Kormanec 2004).

*S. parvulus* have been reported to produce antibiotics like Actinocin, Actinomycin, Borrelidin, Hydroxyectoine and Manumycin (StreptomeDB, www.pharmaceuticalbioinformatics.de/streptomedb). Studies dealing with optimizing such compounds are scarce with the exception of Actinomycin (Foster and Katz 1981; Sousa et al. 2001).However, studies by Genilloud et al. (2011) have shown that inspite of taxonomic relatedness of these strains, the conditions for antibiotic production were strain dependent.

Period of incubation had a profound effect on antibiotic production with maximum zone of inhibition exhibited after 14 days. Further increase in incubation period led to decrease in the antibiotic production which is in accordance to the previous studies reporting that antibiotic production usually occurs in late exponential and stationary phase (El-Nasser et al. 2010; Singh and Rai 2012).

Maximum antibiotic production at 28°C was in agreement to the previous reports indicating that optimal temperature for antibiotic production is usually in the range of 26°C to 35°C exhibited by several *Streptomyces* species (Elliah et al. 2004; Rizk et al. 2007; Mustafa 2009; Ghosh and Prasad 2010; Elleuch et al. 2010; Atta et al. 2011; Mangamuri et al. 2012; Singh and Rai 2012; Vijayakumar et al. 2012; Gunda and Charya 2013).

Antibiotic production in the present study was affected by change in pH of growth medium which is a significant factor affecting nutrient solubility and uptake, enzyme activity, cell membrane morphology by product formation and oxidative reduction reactions (Bajaj et al. 2009; Vijayabharathi et al. 2012). This study found pH 7 as the optimum for antibiotic production and decrease or increase in these values led to complete loss of antibiotic production. Thus the data, is in accordance to the previous reports illustrating pH 7 as optimal for enhancing antibiotic production, exhibited by most *Streptomyces* sp. strains (Elleuch et al. 2010; Singh and Rai 2012; Gunda and Charya 2013). Besides, *S. coelicolar* (Bystrykh et al. 1996), *S. hygroscopicus* D1.5 (Bhattacharya et al. 1998), *S. torulosus* KH-4 (Atta et al. 2010), *S. viridodiastaticus* (El-Nasser et al. 2010), *Streptomyces cheonanensis* (Mangamuri et al. 2012) also stated maximum activity at pH 7. This phenomenon could be attributed to the adaptation of the strain to alkaline soils from which it has been isolated (Rezuanul et al. 2009).

Sources like NaCl had no significant effect on the antibiotic production and were consistent with previous report with respect to neomycin production (Kavitha and Vijayalakshmi 2009).

Antibiotic synthesis is highly dependent on utilization of the preferred carbon sources. The results of this study revealed that maximum antibiotic activity was observed with 0.7% (v/v) glycerol concentration. Glycerol as the better carbon source for enhancing antibiotic production by *Streptomyces* has been reported in the previous studies (Selvin et al. 2009; Elleuch et al. 2010; Singh and Rai 2012; da Silva 2012). According to Shikura et al (2002), when glycerol is used as the precursor, it forms a β-ketoacyl-CoA, a process similar to polyketide biosynthesis where a dihydroxyacetone-type-C_3_ unit is derived from glyce rol to create a β-keto ester leading to a γ-butyrolactone autoregulators which is regarded as *Streptomyces* hormones that trigger the onset of secondary metabolism in general and that of antibiotic production in particular.

Other than the carbon, assimilation of nitrogen source is also crucial for antibiotic production and is regulated by complex mechanisms of glutamate synthetases (Rodríguez-García et al. 2009, Kavitha and Vijayalakshmi 2009; Selvin et al. 2009; Saha et al. 2010; Vijayabharathi et al. 2012; da Silva 2012). Our study revealed 0.03% casein concentration as the optimal for maximum antibiotic production.

Phosphate is also a major factor in antibiotic biosynthesis and expression of phosphate-regulated genes in *Streptomyces* species is modulated by the two-component system PhoR-PhoP (Martin and Demain 1980; Rodríguez-García et al. 2009). Dipotassium hydrogen phosphate (K_2_HPO_4_) is being reported as the most favourable salt for its production. Our results showed 0.25% (w/v) phosphate as optimal for antibiotic production. This data corroborates with the findings of Harold 1966; Kishimoto et al. 1996; Kavitha and Vijayalakshmi 2009; El-Nasser et al. 2010; Mangamuri et al. 2012.

Among the inorganic carbon and nitrogen sources, maximum antibiotic production was obtained with 0.0015% and 0.002% (w/v) concentration of CaCO_3_ and 0.1% (w/v) concentration of potassium nitrate. CaCO_3_ being used as a source of Ca^-2^ enhances antibiotic production and also aids in maintaining the pH of the medium (Hamedi et al. 2004; Basavaraj et al. 2011). Potassium nitrate as superior to other inorganic nitrogen sources has been reported (El-Nasser et al. 2010) and present findings confirms the same.

This study identified a set of optimizing parameters such as a period of 14 days of incubation,with optimum temperature of 28°C and pH of 7 and KusterZ’s modified medium containing glycerol 0.7% (w/v), casein 0.03% (w/v), NaCl 0% (w/v), phosphate 0.25% (w/v), KNO_3_ 0.1% (w/v) and CaCO_3_ 0.0015% (w/v) concentration, that culminated in 11% higher yield of antibiotic production with a mean ZOI of 42 mm against clinical pathogenic strain, *Staphylococcus citreus.* It also highlights the need to screen tropical soil actinobacteria against such rising harmful human pathogen and obtain potential antibiotics that could also serve as targets to MRSA like pathogens.

## Acknowledgements

Authors acknowledge UGC-SAP for providing financial assistance to carry out the work and Dr. Savio Rodrigues, Goa Medical College for providing microbial test pathogens. The first author duly acknowledges the support of UGC-JRF Maulana Azad National Fellowship.

## References

Abbanat D, Maiese W, Greenstein M. Biosynthesis of the pyrroindomycins by Streptomyces rugosporus LL-42D005; Characterization of nutrient requirements. J Antibiot 1999;52:117–126.

Atta HM, Bahobail AS, El-Sehrawi MH. Studies on isolation, classification and phylogenetic characterization of antifungal substance produced by Streptomyces albidoflavus-143. New York Sci J 2011;4:40–53.

Atta HM, Bayoumi R, El-Sehrawi M, Aboshady A, Al-Huminay A. Biotechnological application for producing some antimicrobial agents by actinomycetes isolates from Al-Khurmah Governorate. Euro J Appl Sci 2010;2:98–107.

Badji B, Mostefaoui A, Sabaou N, Lebrihi A, Mathieu F, Seguin E, Tillequin F. Isolation and partial characterization of antimicrobial compounds from a new strain Nonomuraea sp. NM94. J Ind Microbiol Biotechnol 2007;34:403–412.

Bajaj IB, Lele SS, Singhal RS. A statistical approach to optimization of fermentative production of poly(c-glutamic acid) from Bacillus licheniformis NCIM 2324. Bioresour Technol 2009;100:826–832.

Battacharyya BK, Pal SC, Sen SK. Antibiotic production by Streptomyces hygroscopicus D1.5: Cultural effect. Rev Microbiol 1998;29:49–52.

Berdy J. Bioactive microbial metabolites. J Antibiot 2005;58:1–26.

Bystrykh LV, Fernander-Moreno MA, Herremo JK, Malportida F, Hopwood DA, Dijkhuizenen L. Production of actinorhodin-related blue pigments by Streptomyces coelicolor. J Bacteriol 1996;178:2238–2244.

Chambers HF. The changing epidemiology of Staphylococcus aureus? Emerg Infect Dis 2001;7:178–182.

Chhabra SR, Keasling JD. The biological basis | metabolic design and control for production in prokaryotes. In: Moo-Young M, editors. Comprehensive Biotechnology. Elsevier; 2011. p. 243–255.

Cross T, Goodfellow M. Taxonomy and classification of the actinomycetes. In: Sykes G, Skinner FA, editors. Actinomycetales: Characteristics and Practical Importance. New York: Academic Press; 1973. p. 11–91.

da Silva IR, Martins MK, Carvalho CM, de Azevedo JL, de Lima Procopio RE. The Effect of Varying Culture Conditions on the Production of Antibiotics by Streptomyces spp., Isolated from the Amazonian Soil. Ferment Technol 2012;1:2–5.

E. Darabpour, Ardakani MR, Motamedi H, Ronagh MT, Najafzadeh H. Purification and optimization of production conditions of a marine-derived antibiotic and ultra-structural study on the effect of this antibiotic against MRSA. Eur Rev Med Pharmacol Sci 2012;16:157–165.

Demain AL, Sanchez S. Microbial drug discovery: 80 years of progress. J Antibiot 2009;62:5–16.

Devillers J, Steiman R, Seigle MF. The usefulness of the agar-well diffusion method for assessing chemical toxicity to bacteria and fungi. Chemosphere 1989;19:1693–1700.

Dilsen N, Demiroglu C, Ulagay I. A case of arthritis caused by Staphylococcus citreus. Turk Tip Cemiy Mecm 1961;27:275–81.

Elleuch L, Shaaban M, Smaoui S, Mellouli L, Karray-Rebai I, Fourati-Ben Fguira L, Shaaban KA, Laatsch H. Bioactive Secondary Metabolites from a New Terrestrial Streptomyces sp. TN262. Appl Biochem Biotechnol 2010;162:579–593.

Elliah P, Srinivasulu B, Adinarayana K. Optimization studies on Neomycin production by a mutant strain of Streptomyces marinensis in soild state fermentation process. Biochem 2004;39:529–534.

El-Nasser NHA, Helmy SM, Ali AM, Keera AA, Rifaat HM. Production, purification and characterization of the antimicrobial substances from Streptomyces viridodiastaticus (Nrc1). Can J Pure Appl Sci 2010;4:1045–1051.

Felsenstein J. Confidence limits on phylogenies: an approach using the bootstrap. Evolution 1985;39:783–791.

Fischbach MA, Walsh CT. Antibiotics for emerging pathogens. Science 2009;325:1089–2093.

Genilloud O, Gonzalez I, Salazar O, Martin J, Tormo JR, Vicente F. Current approaches to exploit actinomycetes as a source of novel natural products. J Ind Microbiol Biotechnol 2011;38:375–389.

Ghosh UK, Prasad B. Optimization of carbon, nitrogen sources and temperature for hyper growth of antibiotic producing strain Streptomyces kanamyceticus MTCC 324. Bioscan 2010;5:157–158.

Gunda MM, Charya MAS. Physiological factors influencing the production of antibacterial substance by fresh water actinobacteria. J Recent Adv Appl Sci 2013;28:55–62.

Hamedi J, Malekzadeh F, Saghafi-nia AE. Enhancing of erythromycin production by Saccharopolyspora erythraea with common and uncommon oils. J Ind Microbiol Biotechnol 2004;31:447–756.

Harold FM. Inorganic polyphosphates in biology: structure, metabolism and function. Bacteriol Rev 1996;30:772.

Hasegawa M, Kishino H, Yano T. Dating of the human-ape splitting by a molecular clock of mitochondrial DNA. J Mol Evol 1985;22:160–174.

Kavitha A, Vijayalakshmi M. Cultural parameters affecting the production of bioactive metabolites by Nocardia levis MK-VL_113. J Appl Sci Res 2009;5:2138–2147.

Kishimoto K, Park YS, Okabe M, Akiyama S. Effect of phosphate ion on Mildiomycin production by Streptoverticillium rimofaciens. J Antibiot 1996;49:775–780.

Kuster E, Williams S. Selective media for isolation of Streptomycetes. Nature 1964;202:928–929.

Locci R. Streptomycetes and Related Genera. In: Goodfellow M, Williams ST, Mordarski M, editors. Bergey’s Manual of Systematic Bacteriology. New York: Academic Press; 1989. p. 1–32.

Luthra U, Dubey RC. Medium optimization of lipstatin from Streptomyces toxytricini ATCC 19813 by shake flask study. Int J Microbiol Res 2012;4:266–269.

Mangamuri UK, Poda S, Naragani K, Muvva V. Influence of Cultural Conditions for Improved Production of Bioactive Metabolites by Streptomyces cheonanensis VUK-A Isolated from Coringa Mangrove Ecosystem. Curr Trends Biotechnol Pharm 2012;6:99–111.

Martin JF, Demain AL.Control of antibiotic biosynthesis. Microbiol Rev 1980;44:230–251.

Mustafa O. Antifungal and antibacterial compounds from Streptomyces strains. Afri J Biotechnol 2009;8:3007–3017.

Nanjwade BK, Chandrashekhara S, Goudanavar PS, Shamarez AM, Manvi FV. Production of antibiotics from soil-isolated actinomycetes and evaluation of their antimicrobial activities. Trop J Pharma Res 2010;9:373–377.

Olmos E, Mehmood N, Husein LH, Goergen J-L, Fick M, Delaunay S. Effects of bioreactor hydrodynamics on the physiology of Streptomyces. Bioproc Biosyst Eng 2013;36:259–272.

Panda N, Nandi S, Chakraborty T. Isolation, yield optimization and characterization of bioactive compounds from soil bacteria utilizing HPLC and Mass Spectra. Asian J Biomed Pharma Sci 2011;1:01–07.

Pinner M, Voldrich M. Derivation of Staphylococcus albus, citreus and roseus from Staphylococcus aureus. J Infect Dis 1932;50:185–202.

Rezuanul IMd, Jeong YT, Ryu YJ, Song CH, Lee YS. Isolation, identification and optimal culture conditions of Streptomyces albidoflavus C247 producing antifungal agents against Rhizoctonia solani AG2-2. Mycobiol 2009;37:114–120.

Rizk M, Tahany AM, Hanaa M. Factors affecting growth and antifungal activity of some Streptomyces species against Candida albicans. Int J Food Agric Environ 2007;5:446–449.

Rodríguez-García A, Sola-Landa A, Apel K, Santos-Beneit F, Martin J F. Phosphate control over nitrogen metabolism in Streptomyces coelicolor: direct and indirect negative control of glnR, glnA, glnII and amtB expression by the response regulator PhoP. Nucl Acids Res 2009;37:3230–3242.

Saha MR, Ripa FA, Islam MZ, Khondkar P. Optimization of conditions and in vitro antibacterial activity of secondary metabolite isolated from Streptomyces sp. MNK7. J Appl Sci Res 2010;6:453–459.

Selvin J, Shanmughapriya S, Gandhimathi R, Seghal Kiran G, Rajeetha Ravji T, Natarajaseenivasan K, Hema TA. Optimization and production of novel antimicrobial agents from sponge associated marine actinomycetes Nocardiopsis dassonvillei MAD08. Appl Microbiol Biotechnol 2009;83:435–445.

Sevcikova B, Kormanec J. Differential production of two antibiotics of Streptomyces coelicolor A3(2), actinorhodin and undecylprodigiosin, upon salt stress conditions. Arch Microbiol 2004;181:384–389.

Shikura N, Yamamura J, Nihira T. barS1, a gene for biosynthesis of a γ-Butyrolactone autoregulator, a microbial signaling molecule eliciting antibiotic production in Streptomyces species. J Bacteriol 2002;184:5151–5157.

Singh N, Rai V. Optimization of cultural parameters for antifungal and antibacterial metabolite from microbial isolate; Streptomyces rimosus MTCC 10792 from soil of Chhattisgarh. Int J Pharm Pharma Sci 2012;4:20–12.

Sousa M de F V de Q, Lopes C E, Júnior NP. A Chemically Defined Medium for Production of Actinomycin D by Streptomyces parvulus. Braz Arch Biol Technol 2001;44:227–231.

Sujatha P, Bapi Raju KVVSN, Ramana T. Studies on a new marine Streptomycete BT-408 producing polyketide antibiotic SBR-22 effective against methicillin resistant Staphylococcus aureus. Microbiol Res 2005;160:119–126.

Tamura K, Peterson D, Peterson N, Stecher G, Nei M, Kumar S. MEGA5: Molecular Evolutionary Genetics Analysis using Maximum Likelihood, Evolutionary Distance, and Maximum Parsimony Methods. Mol Bio Evol 2011;28:2731–2739.

Velho-Pereira S, Kamat MN. Antimicrobial Screening of Actinobacteria using a Modified Cross-Streak Method. Ind J Pharm Sci 2011;73:223–228.

Velho-Pereira S, Kamat NM. A novel baiting technique for diversity assessment of soil actinobacteria in a laboratory microcosm. Int J Biotech & Biosci 2012;2:217–220.

Vijayabharathi R, Devi PB, Sathyabama S, Bruheim P, Priyadarisini VB. Optimization of resistomycin production purified from Streptomyces aurantiacus AAA5 using response surface methodology. J Biochem Tech 2012;3:402–408.

Vijayakumar R, Panneerselvam K, Muthukumar C, Thajuddin N, Panneerselvam A, Saravanamuthu R. Optimization of Antimicrobial Production by a Marine Actinomycete Streptomyces afghaniensis VPTS3-1 Isolated from Palk Strait, East Coast of India. Ind J Microbiol 2012;52:230–239.

Visalakchi S, Muthumary J. Antimicrobial activity of the new endophytic Monodictys castaneae SVJM139 pigment and its optimization. Afr J Microbiol Res 2009;3:550–556.

Watve MG, Tickoo R, Jog MM, Behole BD. How many antibiotics are produced by the Genus Streptomyces ? Arch Microbiol 2001;176:386–390.

Williams ST, Davies FL. Use of a scanning electron microscope for the examination of actinomycetes. Gen Microbiol 1967;48:171–177.

Yarbrough GG, Taylor DP, Rowlands RT, Crawford MS, Lasure LL. Screening microbial metabolites for new drugs theoretical and practical issues. J Antibiot 1993;46:535–544.

